# Interpretable Biological Sequence Clustering with *i*Clust

**DOI:** 10.64898/2026.04.13.718335

**Authors:** Simeng Zhang, Xinying Liu, Jun Lou, Mudi Jiang, Zengyou He

**Author notes:** Contributing authors.

## Abstract

Biological sequence clustering is a fundamental problem in bioinformatics, yet most existing methods mainly optimize clustering quality or efficiency while offering limited insight into why sequences are grouped together. This restricts their usefulness in downstream analysis, where representative sequences and clear cluster boundaries are often needed. To address this issue, we propose *i*Clust, an interpretable clustering method that characterizes each cluster by a representative prototype and an adaptive radius. By adapting to local sequence structure rather than relying on a single global threshold, *i*Clust produces clusters that are both meaningful and explainable. A final consolidation step further reduces tiny fragments and improves structural stability. Experiments on simulated and real biological sequence datasets show that *i*Clust achieves competitive clustering performance while providing clearer cluster-level explanations than conventional threshold-based methods. In addition to its empirical impact as a practical clustering method for biological sequences, this article opens up new avenues for developing biological sequence clustering approaches from the viewpoint of interpretable machine learning.

## 1 Introduction

The rapid expansion of biological sequence collections has made scalable sequence clustering a central task in bioinformatics. Modern clustering tools are increasingly expected to process millions of sequences [1] [2], and methodological development has therefore focused largely on computational efficiency and scalability. However, real biological sequence datasets are rarely clean: they are often affected by duplicate sequences [3], sequencing errors, low-abundance noise, and contamination [4], while also exhibiting complex underlying data distributions. Under such conditions, efficiency alone is not enough. For both researchers and downstream applications, it is equally important to understand why sequences are grouped together or assigned to different clusters, and what kind of support the resulting clustering can provide for subsequent analysis. Yet interpretability in biological sequence clustering remains a critical but still insufficiently developed aspect. Therefore, interpretable clustering for biological sequence data is an important research direction in urgent need of further development.

A variety of well-established approaches have been employed for biological sequence clustering, including hierarchical clustering, graph-based clustering, model-based approaches, and heuristic greedy incremental methods [5]. Among them, the most widely used tools for large-scale analysis are efficiency-oriented greedy strategies based on global similarity thresholds [6], such as CD-HIT [7], VSEARCH [8], and MMseqs2 [9]. These methods are attractive because they can rapidly construct clusters through predefined thresholds and heuristic representative selection, making them highly practical for large-scale sequence analysis. However, their widespread use has also reinforced a tendency to evaluate clustering algorithms mainly in terms of speed and scalability, while paying relatively less attention to whether the resulting clusters are interpretable or whether the clustering mechanism can recover the true and diverse structure of biological sequence data.

For biological sequence clustering, interpretability is not merely an optional add-on, but an important factor that directly affects the credibility of clustering results. Cluster assignments often influence downstream analyses, such as operational taxonomic unit identification and redundancy removal [10]. At the same time, these tasks usually lack reliable ground-truth [11] or unified references, making it difficult for researchers to rely solely on external evaluation metrics to determine whether the clustering results truly reflect the underlying biological structure. In this context, if we cannot explain why sequences are assigned to a particular cluster and how each cluster boundary is formed, then the stability, reliability, and biological relevance of identified clusters become difficult to assess. This issue is especially pronounced in biological sequence data, which often involves a large number of clusters [12] and highly uneven cluster sizes [3] [13]. Therefore, biological sequence clustering must not only produce sequence partitions but also provide interpretable evidence that supports the validity of the results.

Recent research efforts on interpretable clustering [14] have proposed explanatory frameworks based on rules, decision trees, and prototype representations. However, most of these paradigms are better suited to structured data with explicit features or relatively low-dimensional representations, and therefore do not transfer naturally to biological sequence data. Biological sequences are discrete strings whose relationships are more naturally characterized by edit distance, alignment, or homology than by directly readable numerical features [15]. Moreover, biological sequence clustering often produces a large number of clusters, making rule-based or tree-based explanations increasingly burdensome as the number of clusters grows [16]. In this context, prototype-based explanation [17] through representative sequences provides a more natural and practical form of interpretability. A prototype is an individual that serves as a representative example of its cluster and is defined as the one with the smallest dissimilarity to the other individuals within that cluster. This prototype-based view is well suited to biological sequence analysis, as it can be integrated with downstream workflows such as sequence alignment, database retrieval, and functional annotation.

In fact, many classic biological sequence clustering algorithms [7][8][9][18][19][20] already adopt a prototype-like idea by retaining one representative sequence for each cluster and using it to recruit new members. However, in existing methods, representative sequences are usually selected by heuristic criteria such as sequence length, abundance, or input order [5], with the primary goal of improving efficiency rather than constructing interpretable cluster representations. As a result, the chosen representative may not lie near the center of the cluster, nor may it best summarize the cluster members. In addition, defining all clusters with a single global similarity threshold makes it difficult to accommodate local differences in density and variation, often leading to over-splitting in some regions and improper merging in others [21]. Existing methods therefore indicate which sequence serves as the representative, but they often fail to explain where the effective boundary of the cluster lies and why its members are represented by that sequence.

Motivated by these limitations, we propose *i*Clust, an interpretable biological sequence clustering method based on prototype and adaptive-radius representation. The *i*Clust method estimates local radius from local neighborhoods and iteratively refines both the prototype and the radius of each cluster, so that cluster centers and boundaries are characterized jointly rather than by a representative sequence alone. This representation provides each cluster with both a more informative prototype and an interpretable range of influence, thereby improving the interpretability and reliability of clustering results while maintaining strong clustering performance.

Empirical results show that *i*Clust achieves notable advantages over widely used biological sequence clustering methods in terms of interpretability. Across datasets, the representatives learned by *i*Clust remain much closer to the intrinsic centers of clusters. At the same time, *i*Clust learns cluster-specific boundaries that are substantially more compact. Importantly, these gains are not obtained by over-fragmenting the data. On real datasets, baselines typically produce about 4 times to more than 8 times as many clusters as the underlying structure, whereas *i*Clust remains much closer to the expected cluster granularity. Overall, we not only introduce an interpretable clustering method for biological sequences, but also open the door of a new research direction in biological sequence analysis.

## 2 Results

### 2.1 Overview of the *i*Clust algorithm

To ensure both clustering performance and result interpretability in biological sequence clustering, we propose *i*Clust, whose core idea is to use prototype-radius representation as a unified criterion for cluster interpretation: the prototype represents the cluster center, and the radius characterizes the range covered by the cluster, so that each cluster can not only be identified but also clearly explained. Fig. 1 shows the overall workflow of *i*Clust. Instead of generating the final clustering directly from a single global threshold, the method starts from local scale estimation and then proceeds through several stages, including micro-cluster initialization, iterative prototype-radius refinement, global reassignment, pre-merge cleanup, and cluster merging, ultimately outputting interpretable clustering results with clearly defined centers and boundaries.

**Fig. 1.**
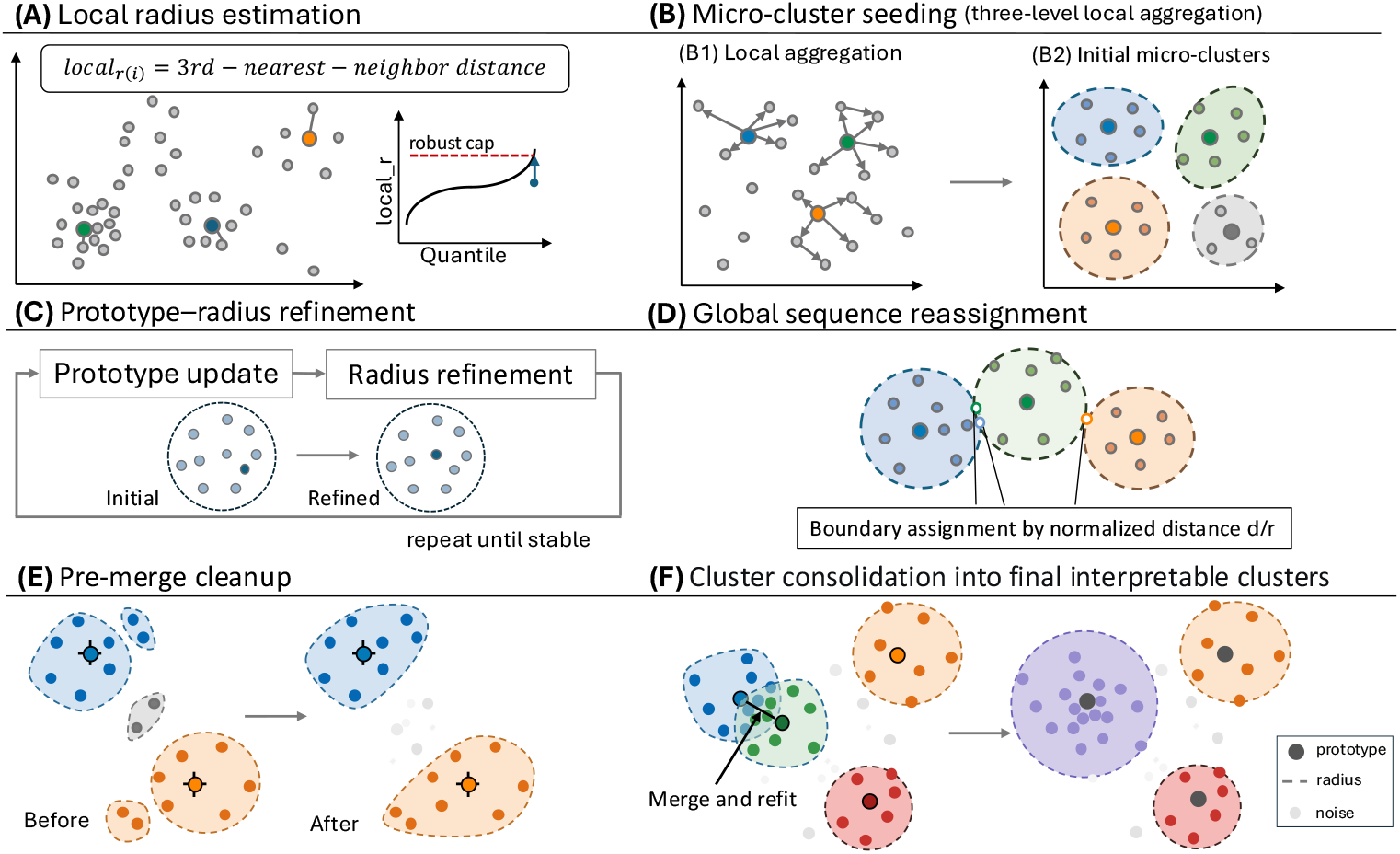
Overview of the *i*Clust algorithm. In **(A)**, the blue, orange, and green sequences are located in regions with different local densities and therefore correspond to local radii of different magnitudes. **(B)** illustrates the aggregation process based on local radii. Sequences are first sorted by their local radii, and the sequence with the smallest radius is selected as the seed. Centered on this seed, the algorithm traverses its neighbor list and assigns as members all unassigned sequences whose distances fall within the seed’s local radius. To ensure sufficient recall, the neighbors of newly added members are further explored to finally obtain a set of small and compact micro-clusters. In **(C)**, these micro-clusters continuously update their prototypes and radii, allowing the initially coarse cluster representations to gradually converge to a more stable prototype-radius structure. In **(D)**, after most sequences have acquired stable assignments, a small number of boundary sequences located near neighboring clusters are reassigned to more suitable clusters according to normalized distance. shows the cleanup step before the final consolidation. Two very small clusters composed of blue or orange sequences are absorbed because they fall within the prototype coverage radius of nearby large clusters. By contrast, the two gray sequences in the middle are dissolved as noise. In **(F)**, the blue and green clusters are merged because their prototypes are sufficiently close and they share a subset of sequences. The prototype and radius are then recalculated, yielding the purple cluster on the right. In the final result, each cluster is jointly described by a prototype and a radius, where the prototype represents the cluster center, the dashed boundary indicates its coverage range, and sequences outside the boundary are retained as noise.

Specifically, *i*Clust first estimates an initial local radius for each sequence, as shown in Fig. 1(A). We use the distance from a sequence to its third nearest neighbor as the initial measure of local radius, so as to capture differences in local density across regions. In this way, high-density regions are associated with smaller local radii, whereas sparse regions are allowed larger local radii. To prevent extreme outliers from producing unreasonably large initial radii and thereby absorbing all samples, we further apply a 99th-percentile truncation to the local radii of all sequences, providing a more stable basis for the subsequent clustering process.

On this basis, *i*Clust uses the local radii obtained in the first step to construct initial micro-clusters. To ensure efficiency, a three-level local aggregation strategy is adopted to rapidly group neighboring sequences, as shown in Fig. 1(B). Rather than directly pursuing the final global partition, this step first forms a set of small and compact candidate clusters from local neighborhoods with strong internal consistency. This design serves two purposes. First, it preserves local structure under complex data distributions and helps avoid premature incorrect merging. Second, it provides a more stable starting point for subsequent prototype and radius learning, so that cluster interpretation is built on a locally reliable foundation from the outset.

Subsequently, *i*Clust iteratively updates the prototype and radius within each micro-cluster, as shown in Fig. 1(C). During this process, the prototype is gradually adjusted toward the position with the smallest overall distance to the rest members of the cluster, while the radius is progressively refined according to recall and precision so as to describe the actual extent of the cluster more accurately. Through this procedure, the representation of the cluster center and boundary is optimized jointly, resulting in a stable prototype-radius representation. Unlike traditional methods that retain only a heuristic representative sequence, this step allows both the cluster representative and its effective coverage to be modeled simultaneously.

After the prototype and radius of each cluster have become stable, *i*Clust performs a global reassignment step (Fig. 1(D)). At this stage, boundary sequences located in the overlapping regions of neighboring clusters are reassigned to the more appropriate cluster according to the normalized distance. Through this process, the representations learned from local micro-clusters are extended to a consistent partition over all sequences, resulting in clusters that are more complete and stable, while also ensuring that sequence assignments remain consistent with the learned prototype-radius representation.

Before producing the final output, *i*Clust further performs a cleanup and consolidation step (Fig. 1(E)–(F)). Among the previously obtained stable micro-clusters, a small number of clusters may be extremely small or fragmented, containing only a few sequences. These clusters may correspond to noise, variants, or boundary components of a larger cluster. From the perspective of interpretability, such tiny clusters require separate handling: they may either be dissolved into noise or, under appropriate conditions, merged into a suitable larger cluster. After these small clusters are processed, neighboring clusters with clear overlap can be further merged, and the prototype and radius of each newly merged cluster are then recalculated. In this way, *i*Clust produces a clustering structure that is more compact and easier to interpret. Each final cluster is jointly described by a prototype and a radius: the prototype represents the cluster center, and the radius defines its effective boundary, while sequences outside the boundary are retained as noise. Therefore, the output of *i*Clust includes not only cluster labels, but also intuitive explanations of cluster centers, cluster boundaries, and sequence assignments.

### 2.2 Experimental setup

To systematically evaluate the effectiveness, robustness, and interpretability of *i*Clust, we conducted experiments on both synthetic datasets and real sequencing datasets. We first used simulated data, in which within-cluster variation, between-cluster boundaries, noise injection, and abundance distributions could be precisely controlled, to verify whether the underlying mechanism of the algorithm behaves as expected. We then evaluated the clustering performance, interpretability, and practical applicability of *i*Clust on real viral sequence data and real 16S amplicon data. Finally, we further examined its scalability on a larger real 16S dataset. Following this design, four datasets with distinct roles were used in this study: the Zymo simulated datasets for mechanism validation, the viral (Influenza A) dataset [22] and the mothur MiSeq SOP V4 dataset [23] for evaluation in real-label scenarios, and the ATCC MSA-1003 full-length 16S dataset for large-scale practical validation.

#### 2.2.1 Datasets

A total of four datasets were used in this study to evaluate the clustering performance and interpretability of *i*Clust from different perspectives. Details of these datasets are summarized in Table 1.

**Table 1.** THE MAIN CHARACTERISTICS OF DATASETS.

| Dataset | #Sequences | #Classes | Average length |
| --- | --- | --- | --- |
| Zymo simulated datasets | 5000 | 8 | 465 |
| Influenza A | 949 | 5 | 1409 |
| mothur MiSeq SOP V4 dataset | 4569 | 21 | 253 |
| ATCC MSA-1003 full-length 16S dataset | 13306 | 20 | 1496 |

The Zymo simulated datasets were used for controlled experimental validation. Reference V3–V4 16S amplicon sequences were generated in silico from the ZymoBIOMICS Microbial Community Standard reference set (Version 3), obtained from the official repository (https://zymo-files.s3.amazonaws.com/BioPool/ZymoBIOMICS.STD.refseq.v3.zip). To systematically examine the ability of *i*Clust to adapt to different levels of complex clustering structures, we designed three mutationrate settings, denoted as easy-, medium-, and hard-mode scenarios (Table 2). In addition, two targeted simulation settings were constructed. The first was a noise-injection scenario, in which the far-noise mutation rate was set to 0.15 and the actual injected noise proportion was 5%, to evaluate the ability of the algorithm to identify and reject abnormal sequences far from the true cluster boundaries. The second was an imbalanced scenario designed to mimic the long-tailed abundance distributions commonly observed in real sequencing data. In this setting, *Bacillus subtilis, Enterococcus faecalis*, and *Escherichia coli* together accounted for approximately 70% of all sequences, while each of the remaining five species accounted for 6.00%. This dataset was used to examine whether *i*Clust can maintain stable clustering quality and interpretable outputs under highly imbalanced cluster sizes.

**Table 2.**
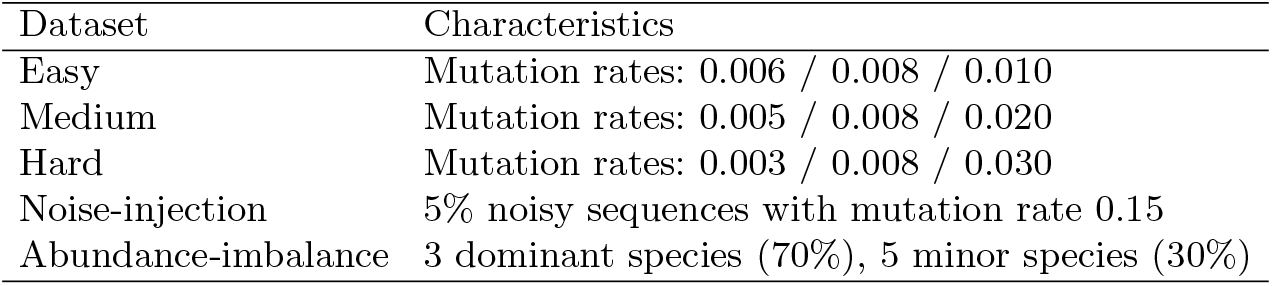
SIMULATION SETTING OF THE ZYMO SIMULATED DATASETS.

The viral dataset was used to evaluate the performance of *i*Clust in a real-world setting. It contains 949 viral sequences from five subtypes, with a relatively balanced class distribution: H7N3 (193), H1N1 (191), H7N9 (190), H5N1 (188), and H2N2 (187). The mothur MiSeq SOP V4 dataset was used as a benchmark for standard 16S amplicon clustering. Compared with the simulated data, these two datasets exhibit more complex within-class heterogeneity and between-class similarity, making them more suitable for examining whether *i*Clust can still produce high-purity, stable, and representative clusters with interpretable prototypes and boundaries under real and heterogeneous conditions. In addition, the ATCC MSA-1003 full-length 16S dataset (https://www.atcc.org/products/msa-1003) was used to assess the applicability and scalability of *i*Clust on a larger real 16S dataset. Taken together, these four datasets play complementary roles, spanning controlled simulation, real-world heterogeneity, standard 16S benchmarking, and larger-scale validation.

#### 2.2.2 Baselines

To comprehensively evaluate the clustering performance and interpretability of *i*Clust, we selected three representative biological sequence clustering methods as baselines: CD-HIT [7], VSEARCH [8], and Clusterize [1]. CD-HIT and VSEARCH are among the most commonly used baseline methods for biological sequence dereplication and rapid clustering. Both are well established, computationally efficient, and widely adopted in practice. Compared with traditional fixed-threshold clustering methods, Clusterize places greater emphasis on improving clustering accuracy and scalability through relatedness sorting, making it an important reference for assessing whether *i*Clust can maintain good clustering quality while also providing interpretable outputs. Together, these three methods cover both widely used mainstream approaches and more recent developments, allowing the performance of *i*Clust to be examined from multiple perspectives. To ensure a fair comparison, all methods were run on the same datasets under a unified evaluation framework.

#### 2.2.3 Evaluation metrics

In this study, all methods were evaluated from two perspectives: clustering quality and interpretability quality. The former measures the consistency between the predicted clusters and the ground-truth labels. The latter evaluates whether the representative sequence has good within-cluster representativeness, whether the cluster radius can effectively characterize the within-cluster boundary, and whether the clustering results remain stable in incremental data scenarios.

##### (1) Interpretability quality

Given that the core output of *i*Clust includes not only the clustering partition itself, but also a representative prototype and an adaptive radius for each cluster, we evaluate interpretability quality from two aspects: the effectiveness of prototype–radius explanations in the static setting, and the applicability of the learned prototype–radius structure to newly arriving sequences in the streaming setting.

Given a dataset *X* = {*x*_1_, *x*_2_, …, *x*_*n*_} containing *n* biological sequences, let *Y* = {*y*_1_, *y*_2_, …, *y*_*n*_} denote the corresponding ground-truth labels, where *y*_*i*_ ∈ {1, 2, …, *C*} and *C* is the number of true clusters. Let *Ŷ* = {*ŷ*_1_, *ŷ*_2_, …, *ŷ*_*n*_} denote the clustering labels produced by a given method, where *ŷ*_*i*_ ∈ {1, 2, …, *K*}∪{−1}, *K* is the number of predicted clusters, and −1 indicates that a sample is treated as noise. For each groundtruth cluster *c*, we denote by *S*_*c*_ = {*x*_*i*_ | *y*_*i*_ = *c*} the set of sequences belonging to *c*. Among all predicted clusters, the cluster with the largest overlap with *S*_*c*_ is taken as its matched cluster, denoted by *Ŝ*_*m*(*c*)_, where *m*(*c*) = arg max_1≤*k*≤*K*_ |*S*_*c*_ ∩ *Ŝ*_*k*_ |. Let *p*_*c*_ and *R*_*c*_ denote the prototype and radius of the matched cluster, respectively. Finally, let *d*(·,·) denote the normalized sequence distance used throughout the evaluation. Based on these notations, we define the Average Representation Error (ARE), Best Average Representation Error (BestARE), ARE-Gap, and boundary coverage (Inlier%) as the core metrics for interpretability evaluation.

For the *c*-th ground-truth cluster *S*_*c*_, we first define the Average Representation Error (ARE) as the average distance from all sequences in the cluster to the prototype, that is,

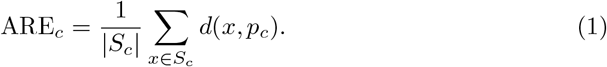

A smaller ARE_*c*_ indicates that the prototype *p*_*c*_ is closer to the center of the sequences within that cluster. However, because different ground-truth clusters may naturally exhibit different levels of internal dispersion, ARE alone is not sufficient to determine whether a prototype is already close enough to the true class center. Therefore, we further enumerate all sequences within each ground-truth cluster as candidate prototypes and define the Best Average Representation Error (BestARE) as:

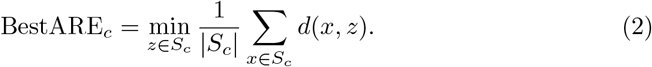

BestARE can be regarded as the theoretical optimal medoid achievable within the ground-truth cluster under the current distance measure. Based on this, we define the ARE-Gap as the ratio of the actual prototype error to the theoretical optimal error:

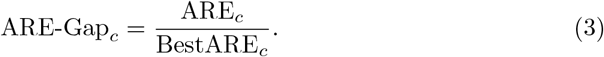

A value of ARE-Gap closer to 1 indicates that the prototype produced by the current method is closer to the one with BestARE within that ground-truth cluster. In contrast, a value substantially greater than 1 suggests that there is still a considerable gap between the selected prototype and the optimal representative.

In addition to prototype representativeness, we further evaluate the effectiveness of the radius boundary. Let *R*_*c*_ denote the radius of the cluster matched to the *c*-th ground-truth class. We then define Inlier% as the proportion of sequences in that ground-truth cluster that fall within the prototype radius, i.e.,

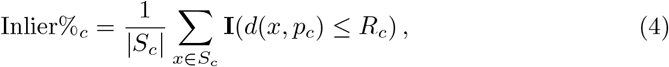

where **I**(·) is the indicator function, which takes the value 1 when the condition inside the parentheses is satisfied and 0 otherwise. A higher value of Inlier%_*c*_ indicates that the explanatory structure defined by prototype *p*_*c*_ and radius *R*_*c*_ is better able to cover the main members of the corresponding ground-truth cluster, and therefore that the boundary semantics of the radius are more effective. Unlike label-consistency metrics alone, Inlier% directly reflects whether the prototype-radius explanation truly covers most sequences in that cluster. Overall, ARE-Gap is mainly used to evaluate the representativeness of the prototype, whereas Inlier% is primarily used to assess the coverage ability of the radius boundary. Together, these two metrics constitute the core criteria in this study for evaluating the effectiveness of the prototype-radius explanation.

Beyond the static evaluation of prototype representativeness and radius boundary validity, we further assess the interpretability of *i*Clust under a streaming setting. The central question is whether the learned prototype–radius explanations remain effective as new sequences are continuously introduced.

Let *D*^(0)^ ⊂ *X* denote the initial training set, and let the model trained on *D*^(0)^ produce prototypes 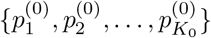 and radii 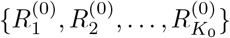. For an incoming batch 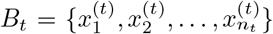, each sequence is assigned to an existing cluster if it falls within at least one learned radius; if multiple clusters satisfy this condition, it is assigned to the one with the smallest normalized score 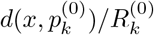. Otherwise, it is rejected and assigned −1.

First, the streaming coverage on batch *B*_*t*_ is defined as:

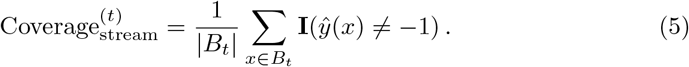

We then define covered-sequence consistency which evaluates the applicability of the learned prototype–radius structure to newly arriving sequences. Let major(*k*) denote the dominant ground-truth label of cluster *k* in the initial model. Then,

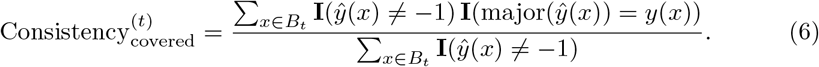

If the learned prototypes and radii are meaningful, then newly absorbed samples should be more likely to receive cluster assignments consistent with their true labels. Therefore, a higher consistency indicates that the learned prototype–radius structure not only covers incoming sequences, but also indirectly supports the validity of the learned explanations. Since this metric is computed only over the absorbed samples, it should be interpreted together with streaming coverage.

##### (2) Clustering quality

We evaluated clustering quality using Adjusted Rand Index (ARI) and Normalized Mutual Information (NMI) to measure the agreement between the predicted cluster labels and the ground-truth labels. In addition, the Silhouette and Davies Bouldin index (DB) were adopted to further assess clustering performance from the perspectives of within-cluster compactness and between-cluster separation. The value ranges and preferred directions of these metrics are summarized in Table 3.

**Table 3.** THE CHARACTERISTICS OF CLUSTERING VALIDITY INDICES.

| CVIs | Range | Optimal directions | Reference |
| --- | --- | --- | --- |
| ARI | $[-1, 1]$ | ↑ | [24] |
| NMI | $[0, 1]$ | ↑ | [25] |
| DB | $[0, +\infty)$ | ↓ | [26] |
| Silhouette | $[-1, 1]$ | ↑ | [27] |

### 2.3 *i*Clust learns representative prototypes and meaningful radii

Fig. 2 systematically evaluates the interpretability of *i*Clust under both static and streaming settings. Overall, *i*Clust shows strong prototype representativeness, boundary coverage, and stable applicability of its explanatory structure under incremental scenarios across different types of datasets. This suggests that the learned prototype-radius explanation is not valid only for a single clustering result, but has broader robustness and general applicability.

**Fig. 2.**
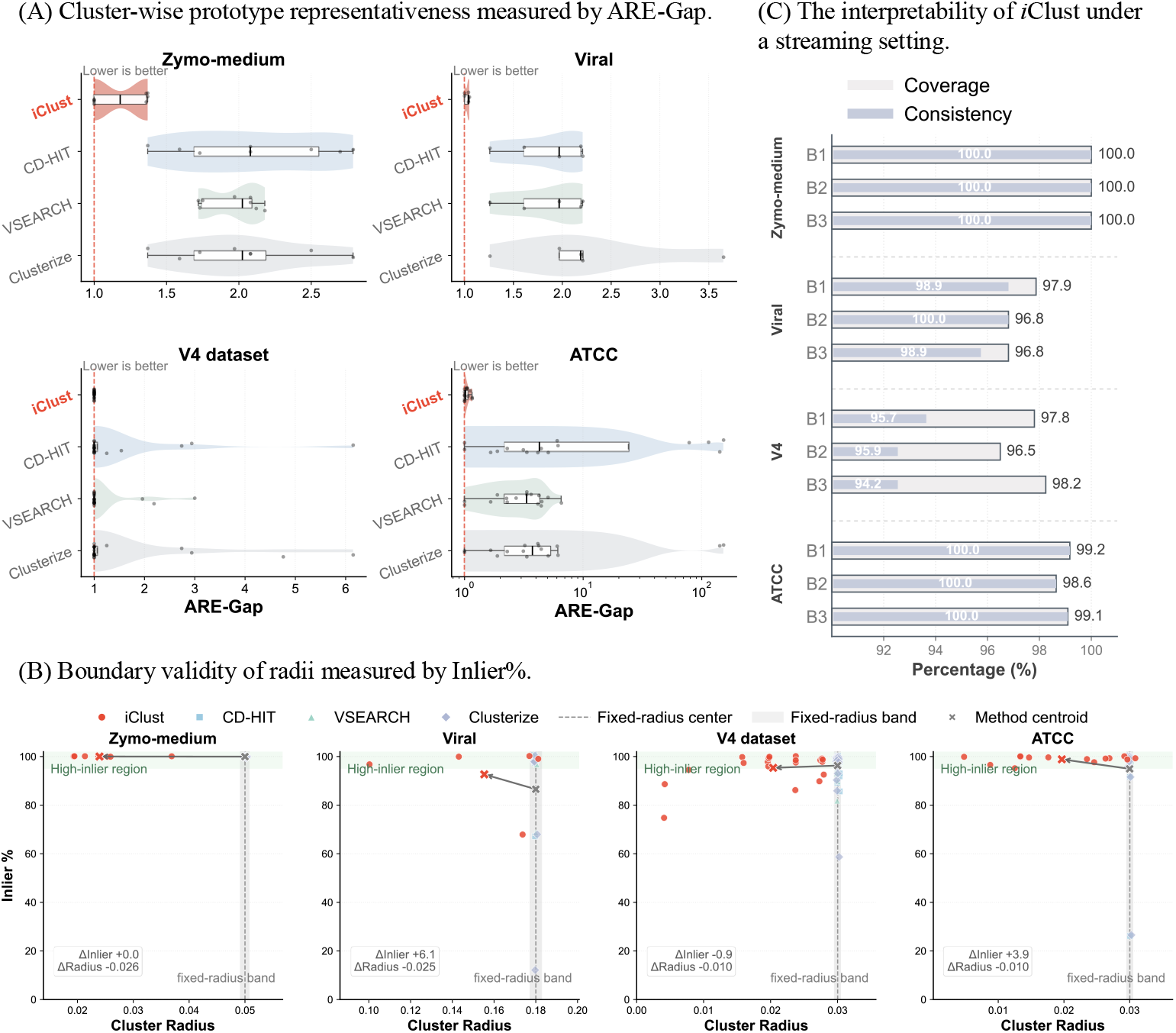
Interpretability of *i*Clust in static and streaming settings. The similarity threshold is kept consistent across all methods for the Zymo simulated dataset, the viral dataset, the V4 dataset, and the ATCC dataset. For *i*Clust, the initial radius is set to 0.05, 0.18, 0.03, and 0.03 for the four datasets, respectively. For the baseline methods, the similarity thresholds are set to 0.95, 0.82, 0.97, and 0.97, respectively (for a more detailed discussion of how similarity thresholds affect the results of the algorithms, see Supplementary section 1).

In Fig. 2(A), we use ARE-Gap to measure prototype representativeness. By definition, an ARE-Gap value closer to 1 indicates that the prototype selected by the algorithm is closer to the most central sequence of the corresponding ground-truth cluster. As shown in Fig. 2(A), *i*Clust achieves the lowest and most stable ARE-Gap across all four datasets, with most clusters distributed tightly around 1. This indicates that the prototypes learned by *i*Clust are generally located near the cluster centers. In contrast, CD-HIT, VSEARCH, and Clusterize show consistently higher ARE-Gap values and noticeably larger variation, suggesting that their selected representative sequences are more likely to deviate from the true centers of the clusters. This difference is particularly evident on the more complex real datasets, namely the V4 and ATCC datasets, where the baseline methods exhibit greater dispersion and substantially inflated ARE-Gap values for some clusters, indicating weaker stability in representative selection. Overall, Fig. 2(A) shows that *i*Clust can maintain strong prototype representativeness across different levels of data complexity, thereby providing a solid basis for subsequent interpretability analysis.

Fig. 2(B) further examines the effectiveness of the learned radius. Here, the horizontal axis represents the radius learned for each cluster, and the vertical axis shows Inlier%, which reflects whether the explanatory structure defined by the prototype as the center and the radius as the boundary can cover the main members of the corresponding ground-truth cluster. As shown in Fig. 2(B), *i*Clust generally achieves higher Inlier% across all four datasets, with most clusters concentrated at high values, in some cases even reaching 100%. This indicates that its adaptive radius can better fit the differences among clusters. By contrast, the baseline methods use a fixed threshold which can be treated as radii, so their results are concentrated along vertical lines and lack the ability to adjust cluster boundaries according to cluster-specific variation. Although a fixed radius can still achieve high Inlier% for some clusters, it performs very poorly for others, with some cases even dropping to under 20%, showing that a single threshold is insufficient to handle the heterogeneity across different clusters. In comparison, the distribution of *i*Clust shows a clear pattern of variable radii with consistently stable coverage: different clusters are allowed to learn different radii, while overall maintaining high Inlier%. This result demonstrates the effectiveness of the learned radius in capturing the boundaries of different clusters.

Fig. 2(C) illustrates the applicability of the learned prototype-radius explanation under a streaming setting, aiming to examine whether it can indeed provide meaningful insight. In this experiment, the model was first trained on 70% of the sequences, and the remaining sequences were then injected into the model in three batches *B*_1_, *B*_2_, and *B*_3_ (each batch contains 10% sequences). We report two metrics, coverage and consistency: coverage measures how many newly arriving sequences can be covered by the original explanatory structure, whereas consistency measures whether the absorbed sequences can be correctly explained by the learned prototype-radius representation.

Overall, all four datasets maintain high coverage and consistency across the three insertion stages, indicating that the explanatory structure learned by *i*Clust is reusable rather than being a one-time and fragile local result. More specifically, on Zymo-medium, both metrics remain close to 1 throughout, showing that under a controlled simulation setting, the prototype-radius explanation continues to hold almost perfectly after incremental insertion. The viral dataset is also highly stable in the first two stages, although coverage shows a slight decline in the third stage, suggesting that newly introduced sequences in real viral data exhibit greater diversity and therefore pose a stronger challenge to the existing boundaries. Nevertheless, consistency remains high, indicating that the absorbed sequences are still assigned to largely appropriate clusters. On the V4 dataset, both coverage and consistency are slightly lower than those of the first two datasets, and consistency further declines in the third stage. This suggests that the real 16S dataset has stronger heterogeneity and more complex boundary structure, although consistency still remains a high level. In comparison, although ATCC is larger in scale and more complex as a real dataset, both coverage and consistency remain at high levels, showing that the explanatory structure of *i*Clust retains good stability even on larger real-world data.

Overall, Fig.2 shows that the interpretability of *i*Clust has two main advantages. First, in the static setting, its prototypes are closer to the representative centers of the ground-truth clusters, and the learned adaptive radii are better able to cover the main body of each cluster, thereby yielding a reliable prototype-radius explanation. Second, in the dynamic setting, this explanatory structure maintains high coverage and consistency as new sequences continue to arrive, indicating that its interpretability does not depend on a single particular partition but retains validity over time. In other words, the interpretability of *i*Clust lies not only in producing explainable results, but also in providing an explanatory structure that remains reusable.

### 2.4 Competitive clustering quality across simulated and real datasets

Fig. 3 compares the clustering quality of *i*Clust and the three baseline methods from four perspectives: ARI, NMI, Coverage, and the ratio of predicted to true cluster numbers (the similarity threshold is set in the same manner as in Fig. 2, for detailed analyses of Silhouette coefficient, and Davies–Bouldin index see the Supplementary section 1). ARI and NMI measure the agreement between clustering results and ground-truth labels, Coverage reflects the proportion of sequences effectively assigned to non-noise clusters, and the predicted/true cluster ratio is used to indicate whether a method shows obvious over-segmentation. Overall, *i*Clust achieves strong label consistency on most datasets and provides a better balance between coverage and a reasonable number of clusters.

**Fig. 3.**
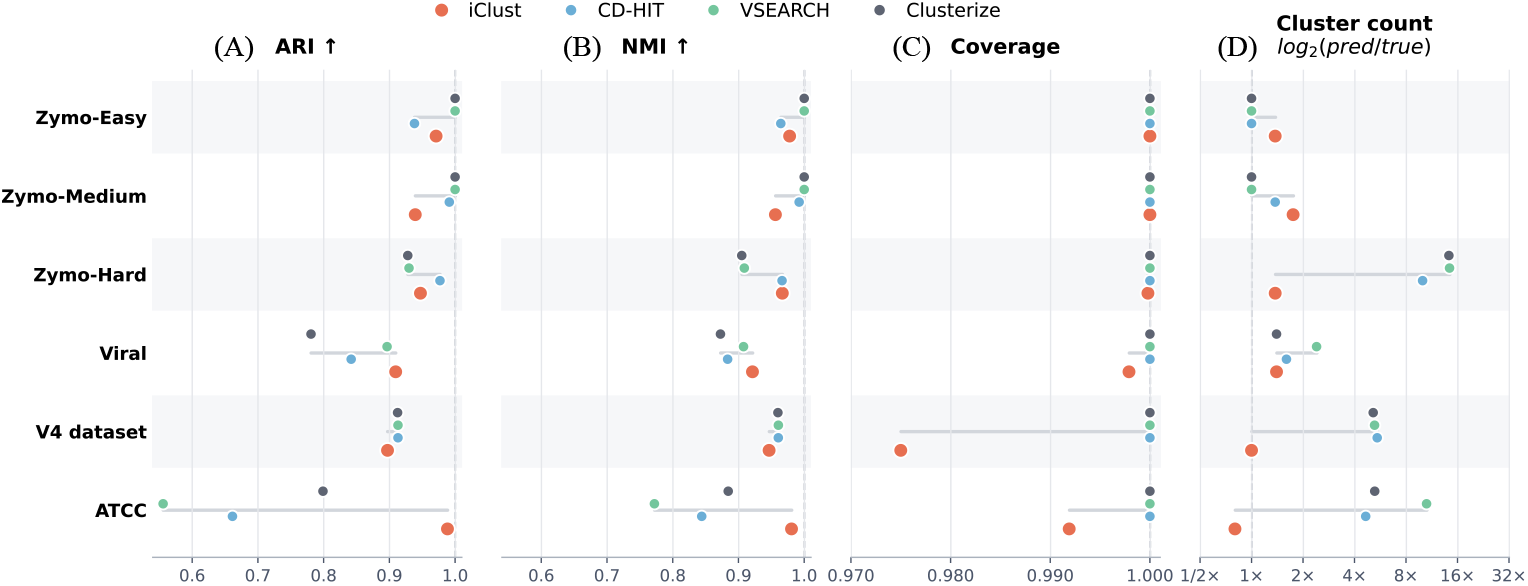
Clustering quality of *i*Clust and baseline methods across datasets.

As shown in Fig. 3 (A) and (B), *i*Clust consistently maintains high ARI and NMI on the three Zymo simulated datasets and also performs stably on the viral dataset. Its advantage becomes even more evident on the more complex ATCC dataset, where both ARI and NMI are clearly higher than those of the three baseline methods, indicating that *i*Clust can still recover cluster structure effectively in real and complex settings. On the V4 benchmark, the ARI and NMI of *i*Clust are slightly lower than those of the best-performing baseline, but the overall difference remains small.

Taken together, Fig. 3(C) and (D) further show that the performance of *i*Clust is not achieved simply by producing more clusters. Although the baseline methods attain Coverage values close to 1 on nearly all datasets, their predicted/true cluster ratios are often much greater than 1 on the more complex datasets, indicating substantial over-segmentation. This is particularly evident on V4 and ATCC, where the baseline methods typically generate far more clusters than the true number of clusters. By contrast, the number of clusters produced by *i*Clust remains much closer to the true cluster number, allowing it to maintain high ARI and NMI while avoiding severe structural fragmentation. It should be noted that the Coverage of iClust on Viral, V4, and ATCC is slightly below 1, mainly because it rejects sequences that cannot be assigned to any prototype-radius structure and treats them as noise, rather than forcing all sequences into clusters.

Overall, Fig. 3 shows that the advantage of *i*Clust lies not only in its strong clustering consistency, but also in its ability to achieve a better overall balance among clustering quality, sequence coverage, and the reasonableness of the resulting cluster structure. This advantage becomes even more pronounced on complex real datasets.

### 2.5 Robustness under noisy and imbalanced settings

As shown in Fig. 4, *i*Clust demonstrates strong robustness in both noisy and long-tailed imbalanced scenarios (details about clustering validity indices and interpretability quality are provided in the Supplementary section 2). As shown in Fig. 4 (A) and (B), on the Zymo noise-injection dataset, *i*Clust can directly identify as noise those sequences that are not accepted by any prototype-radius structure, without requiring additional post-processing. In this experiment, all 240 injected noisy sequences were correctly rejected, with no normal sequences misclassified (FP = 0, FN = 0). In contrast, the initial results of CD-HIT, VSEARCH, and Clusterize produced a large number of fragmented clusters and therefore required post hoc filtering of small clusters to achieve approximate denoising. To ensure a fair comparison, the same minimum cluster size parameter (*minsize* = 3) used in *i*Clust was also applied to the baseline methods, and clusters containing fewer than `minsize` samples were dissolved and treated as noise. The original numbers of clusters produced by CD-HIT, VSEARCH, and Clusterize reached 252, 252, and 260, respectively. After filtering, although the number of clusters decreased, 4, 4, and 5 true sequences were still incorrectly removed by the three methods. These results indicate that the noise handling of *i*Clust does not rely on post-hoc correction, but is instead naturally achieved through its built-in rejection mechanism, allowing it to suppress noise while better preserving the true underlying structure.

**Fig. 4.**
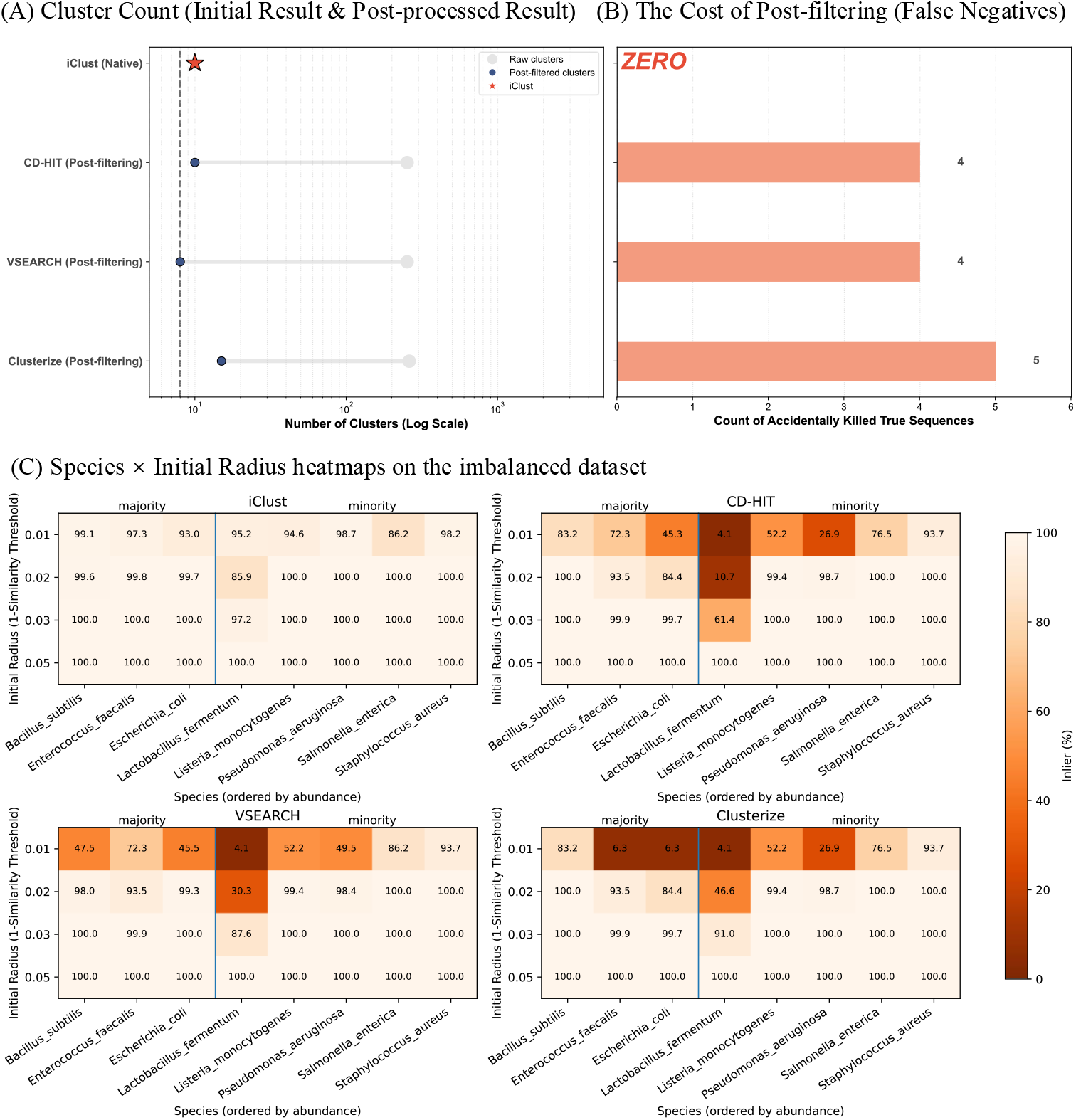
Robustness of *i*Clust and baseline methods under noisy and imbalanced settings. In (A) and (B), the dataset used is the Zymo noise-injection dataset. In this experiment, 240 noise sequences were injected, with a mutation rate of 15%. For *i*Clust, the initial radius was set to 0.05, whereas for the baseline methods, the similarity threshold was uniformly set to 0.95. The dataset used in (C) is the Zymo abundance-imbalance dataset. As described in Table 2, this dataset consists of eight species, among which three species each account for 23.33% of the total and together make up 70%, corresponding to the majority group on the *x*-axis. The remaining five species each account for 6% and together make up 30%, corresponding to the minority group on the *x*-axis. For *i*Clust, the initial radius was set to 0.01, 0.02, 0.03, and 0.05. Correspondingly, for the baseline methods, the similarity thresholds were set to 0.99, 0.98, 0.97, and 0.95, respectively.

On the Zymo abundance-imbalance dataset, *i*Clust shows strong stability. Fig. 4 (C) indicates that, even under relatively strict thresholds, *i*Clust maintained generally high Inlier% values for both majority and minority classes (all above 85%). In contrast, the baseline methods based on a fixed global threshold were more likely to impair tail classes first. For example, under higher similarity thresholds, the Inlier% of *Lactobacillus fermentum* decreased markedly in baseline methods. Overall, the interpretability metric Inlier% proposed in this study shows that, even under class abundance imbalance, *i*Clust can still learn prototypes and radii that match the internal structure of each species. This suggests that *i*Clust not only preserves strong clustering performance but also maintains the effectiveness of its interpretability mechanism.

## 3 Discussion

The *i*Clust method proposed in this study is designed to improve both clustering quality and interpretability in biological sequence clustering. Unlike traditional sequence clustering methods that rely mainly on a fixed global threshold, *i*Clust explains each cluster through a prototype–radius pair, allowing the clustering result not only to reflect the structural relationships among sequences, but also to be interpreted and reused in a more intuitive way. Experimental results across multiple datasets show that *i*Clust can maintain strong clustering performance while also providing cluster-level explanations that are easy to understand. These findings suggest that interpretability and clustering effectiveness are not inherently in conflict, but can be achieved together through an appropriate modeling framework.

The effectiveness of *i*Clust mainly lies in its ability to adapt to the distinctive characteristics of biological sequence data. Such data often show substantial variation in local density, irregular cluster boundaries, and pronounced long-tailed distributions, making it difficult for methods based on a fixed global threshold to handle both dense and sparse regions effectively at the same time. *i*Clust first estimates a local radius for each sequence and thus assigns an adaptive initial radius for aggregation, which reduces the risk of over-fragmentation or excessive merging caused by a single global threshold. It then combines prototype updating with boundary-aware radius refinement, allowing both the cluster center and the cluster boundary to better match the true internal structure of each cluster. Finally, through a conditional merging strategy, neighboring clusters are consolidated in a cautious manner. This stage-wise design enables the method to remain stable under scenarios involving noise, imbalanced distributions, and streaming data insertion.

Despite these strengths, *i*Clust still has several limitations. First, the current method relies on neighborhood candidate construction, distance refinement, and iterative updates of prototypes and radii, and therefore remains more time-consuming overall than one-pass greedy clustering methods based on a fixed threshold. This issue may become more pronounced as dataset size increases, sequence length grows, or cluster boundaries become more complex, because local distance computation, boundary refinement, and cluster merging can become computational bottlenecks. Second, although approximate neighborhood search helps reduce the cost of exhaustive pairwise distance computation to some extent, the current implementation still has limited scalability to very large datasets and is therefore better suited to medium-scale applications or scenarios where interpretability is particularly important. Future work may improve its scalability through more efficient approximate indexing, parallel implementation, block-wise or streaming optimization, and lighter prototype updating strategies. In addition, further reducing parameter sensitivity and extending prototype-radius explanations to more complex settings, such as low-similarity data or hierarchical structures, remain important directions for future research.

## 4 Method

### 4.1 Problem formulation

Given a biological sequence dataset *X* = {*x*_1_, *x*_2_, …, *x*_*n*_}, let *Y* = {*y*_1_, *y*_2_, …, *y*_*n*_} denote the corresponding ground-truth labels, where *y*_*i*_ ∈ {1, 2, …, *C*} and *C* is the number of true clusters. The goal of *i*Clust is to partition the sequences into a set of clusters, with unassigned sequences labeled as noise. Each cluster is represented by a prototype–radius pair, which specifies its center and effective boundary.

Let *d*(*x*_*i*_, *x*_*j*_) denote the distance between two sequences *x*_*i*_ and *x*_*j*_. In this study, we use the normalized Levenshtein distance:

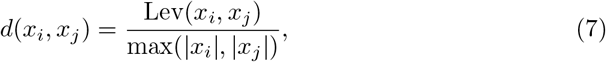

where Lev(·, ·) is the Levenshtein edit distance and |*x*_*i*_| denotes the length of sequence *x*_*i*_.

The output of *i*Clust is a label vector *Ŷ* = {*ŷ*_1_, *ŷ*_2_, …, *ŷ*_*n*_}, where *ŷ*_*i*_ ∈ {1, 2, …, *K*} ∪ {−1}, *K* is the number of final clusters, and *ŷ*_*i*_ = −1 indicates that sequence *x*_*i*_ is treated as noise. For each predicted cluster *C*_*k*_ = {*x*_*i*_ | *ŷ*_*i*_ = *k*}, *i*Clust reports a prototype-radius pair.

- a *prototype p*_*k*_, which serves as the center of cluster *C*_*k*_;
- a *radius R*_*k*_ *>* 0, which defines the effective coverage boundary of cluster *C*_*k*_.

Thus, cluster *C*_*k*_ is characterized by the pair (*p*_*k*_, *R*_*k*_), based on this representation, a sequence *x*_*i*_ can be assigned to cluster *C*_*k*_ only if *d*(*x*_*i*_, *p*_*k*_) ≤ *R*_*k*_. If a sequence can be allocated to multiple clusters, it is assigned to the one with the smallest normalized boundary score:

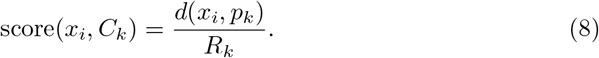

Accordingly, the final assignment rule is defined as:

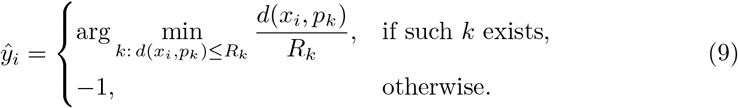

Based on this formulation, each cluster is described by an explicit center–boundary structure, while sequences that cannot be allocated to any cluster are reported as noise.

### 4.2 Implementation of *i*Clust

#### Approximate neighborhood construction and local radius estimation

To initialize cluster formation in a density-adaptive manner, *i*Clust first estimates a local radius for each sequence. Since computing all-pairs distances directly is expensive for large biological sequence datasets, we first construct an approximate neighborhood for each sequence and then compute its precise local radius within that neighborhood. This design is consistent with the overall workflow shown in Fig. 1(A), where local radius estimation serves as the starting point of the method.

For each sequence *x*_*i*_ ∈ *X*, we first generate a compact signature based on its *k*-mer composition and use a banded MinHash–LSH scheme to retrieve a candidate neighbor set 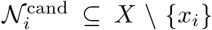 (the parameter details are presented in Section 3 of the Supplementary). In the implementation, these candidates are used only as a high recall filter; the final local radius is still determined from exact normalized Levenshtein distances computed on the retrieved candidates. When the number of retrieved candidates is insufficient, additional sequences are randomly sampled and added to the candidate set. The construction of the neighborhood can also provide a fixed-length candidate list for the subsequent seeding step.

Given the candidate neighbor set for *x*_*i*_, we compute the exact distances 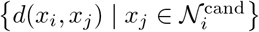, then sort them in a non-decreasing order, and define the initial local radius of *x*_*i*_ as the distance to its *k*-th nearest neighbor:

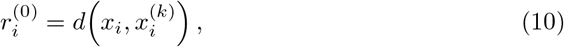

where 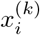 denotes the *k*-th nearest sequence to *x*_*i*_ within the refined candidate set. In this study, we use *k* = 3. Smaller values of 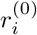 indicate denser local regions, whereas larger values indicate sparser regions.

To reduce the influence of outliers or isolated sequences, *i*Clust further applies a robust upper truncation to the initial local radii. Specifically, the final local radius is obtained by clipping 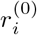 at the 99th percentile of all the initial local radii, with an additional predefined upper bound to prevent a small number of isolated sequences from producing unrealistically large radii:

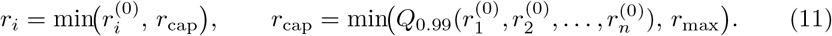

In our implementation, *r*_max_ = 0.05. This truncation improves the stability of subsequent clustering steps.

As a result, each sequence is associated with an adaptive local radius, {*r*_1_, *r*_2_, …, *r*_*n*_}, which serves as the basis for the micro-cluster seeding step described next.

#### Micro-cluster seeding

Given the adaptive local radii estimated above, *i*Clust next performs a microcluster seeding step to obtain conservative initial clusters for subsequent refinement. To reduce over-fragmentation caused by extremely small local radii, we define for each sequence *x*_*i*_ an effective seed radius as:

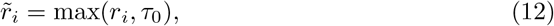

where *r*_*i*_ is the clipped local radius in the previous step and *τ*_0_ is a user-specified initial threshold. Importantly, *τ*_0_ is not used as a hard global clustering threshold. When *r*_*i*_ *< τ*_0_, the seed radius is raised to *τ*_0_ to avoid overly fragmented initialization in very dense regions; when *r*_*i*_ ≥ *τ*_0_, the larger local radius is retained. Therefore, *i*Clust starts clustering around a common initial scale, while still allowing looser local structures to grow according to their own estimated radii.

*i*Clust then sorts all sequences in a non-decreasing order of 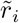 and scans them one by one. Each time an unassigned sequence *x*_*s*_ is encountered, it is selected as a new seed, and its seed radius is fixed as 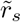 for the current seeding process. Starting from *x*_*s*_, *i*Clust initializes a new micro-cluster and absorbs unassigned sequences according to the seed-centered admissibility condition:

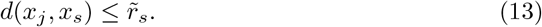

The aggregation is carried out in a local 3-level manner (shown in Fig. 1(B)). First, *i*Clust examines the fixed-length candidate neighbor list returned by the approximate neighborhood construction stage and absorbs all unassigned sequences satisfying Eq. (13). Next, for each newly absorbed member, its candidate neighbors are further explored to retrieve additional admissible sequences that may have been missed in the first pass. Importantly, these newly discovered candidates are still validated against the same seed-centered boundary in Eq. (13), rather than against the radii of intermediate members. Thus, newly added members only expand the search range, but do not redefine the current cluster boundary. To further reduce the number of missed cluster members introduced by approximate neighborhood retrieval, *i*Clust additionally performs a lightweight random check over a small subset of the remaining unassigned sequences. Any sequence satisfying Eq. (13) is also absorbed into the current microcluster. This design improves recall at the seeding stage while keeping the computation efficient.

After 3-level aggregation, the resulting micro-cluster is denoted by 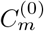. At this stage, *i*Clust assigns it an initial cluster-radius pair:

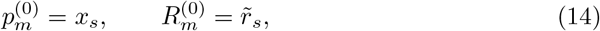

Where 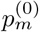 is the initial prototype and 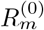 is the initial radius. All members of 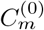 are then removed from the unassigned set, and the procedure continues with the next unassigned sequence in the ordered list.

Repeating this process over all sequences yields a set of initial micro-clusters 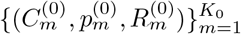, where *K*_0_ is the number of seeded micro-clusters. These initial clusters are intentionally conservative: they provide explicit prototype–radius explanations from the beginning, while leaving prototype updating and radius refinement to the subsequent stage.

#### Prototype update, boundary refinement, and normalized reassignment

To obtain more representative prototypes and more faithful cluster boundaries, *i*Clust further performs an alternating refinement stage. For clarity, Fig. 1(C) and Fig. 1(D) separately illustrate boundary refinement and reassignment, whereas in the actual algorithm they are coupled within the same iterative procedure.

Given the current cluster set 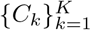, *i*Clust first updates the prototype of each cluster to a medoid-like representative near its center. Specifically, the new prototype is selected from the current cluster members so as to minimize its total distance to other members. Rather than exhaustively evaluating all members, we sample a small candidate subset 𝒜_*k*_ ⊆ *C*_*k*_ and a support subset ℬ_*k*_ ⊆ *C*_*k*_, and define:

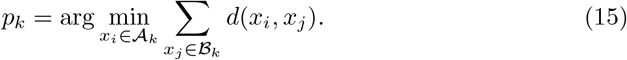

This update keeps the prototype as an actual biological sequence while moving it toward the center of the current cluster.

After the prototype is updated, *i*Clust relearns the cluster radius considering both precision and recall. For cluster *C*_*k*_ with prototype *p*_*k*_, let 𝒫_*k*_ = {*d*(*p*_*k*_, *x*_*i*_) | *x*_*i*_ ∈ *C*_*k*_} denote the set of distances from the prototype to its current members, and let 𝒩_*k*_ = {*d*(*p*_*k*_, *x*_*j*_) | *x*_*j*_ ∉ *C*_*k*_, *x*_*j*_ is sampled from nearby competing clusters} denote a nearby negative set. For any candidate radius *R >* 0, we define:

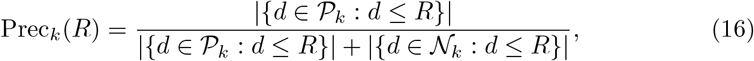

and

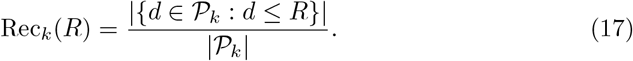

Based on these quantities, the quality of radius *R* is measured by the *F*_*β*_ score:

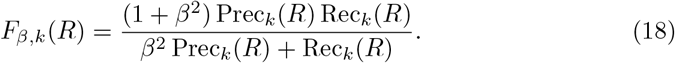

The refined radius is then selected as:

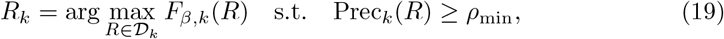

where 𝒟_*k*_ is a finite set of candidate radii 𝒫_*k*_ ∪ 𝒩_*k*_, and *ρ*_min_ is the minimum precision requirement. In this way, the learned radius is encouraged to cover the main body of the cluster while remaining separated from nearby non-members.

Once all prototype–radius pairs (*p*_*k*_, *R*_*k*_) are updated, *i*Clust performs a global reassignment step according to the Eq. (8). Specifically, each sequence is considered admissible only to clusters satisfying *d*(*x*_*i*_, *p*_*k*_) ≤ *R*_*k*_, and if multiple admissible clusters exist, it is reassigned to the one with the smallest normalized score *d*(*x*_*i*_, *p*_*k*_)*/R*_*k*_. Sequences that do not satisfy any admissibility condition remain unassigned and are treated as noise.

The above three operations, prototype update, boundary refinement, and normalized reassignment, are repeated until the cluster configuration becomes stable or the maximum number of iterations is reached. Through this process, the seed-generated micro-clusters are gradually transformed into clusters with more representative prototypes and more effective boundaries, which provides the basis for the subsequent cleanup and consolidation steps.

#### Pre-merge cleanup of tiny fragments

Before cluster consolidation, *i*Clust removes unstable tiny fragments. Sequences in clusters smaller than a predefined minimum size threshold are checked against the currently learned large clusters. If a sequence is admissible to at least one large cluster, it is absorbed into the one with the smallest normalized score *d*(*x*_*i*_, *p*_*k*_)*/R*_*k*_; otherwise, it remains unassigned. Any cluster still below the minimum size threshold is then dissolved. This step removes potentially noisy clusters and yields a cleaner cluster set for the subsequent merge stage.

#### Cluster consolidation into final interpretable clusters

After pre-merge cleanup, *i*Clust further consolidates neighboring clusters into the final interpretable output. Rather than merging clusters solely based on prototype proximity, *i*Clust jointly considers prototype closeness and boundary compatibility.

For two clusters *C*_*a*_ and *C*_*b*_ with prototype–radius pairs (*p*_*a*_, *R*_*a*_) and (*p*_*b*_, *R*_*b*_), *i*Clust first requires their prototypes to be sufficiently close. It then evaluates whether the two clusters are mutually compatible by measuring the acceptance fraction of sampled members across boundaries:

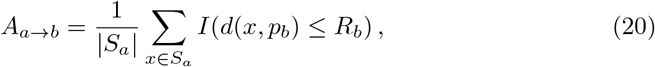

where *S*_*a*_ is a sampled subset of members from *C*_*a*_. The reverse quantity *A*_*b*→*a*_ is defined in the same way. The two clusters are merged only when their prototypes are sufficiently close and the bidirectional acceptance ratio is sufficiently high.

Once a set of clusters is merged, *i*Clust performs a light refit of the merged clusters by re-updating their prototypes and radii and reassigning sequences according to the normalized boundary rule like the refinement stage. This step restores the consistency of the prototype–radius explanation after consolidation.

Finally, any remaining tiny fragments are processed in the same manner as in the pre-merge cleanup step. Through this process, *i*Clust produces a more compact and stable set of final clusters, each described by an interpretable prototype–radius pair.

## Code availability

*i*Clust and the data underlying this manuscript are available at: https://github.com/Helen1momo/iclust.

## Acknowledgements

This work has been supported by the Natural Science Foundation of China under Grant No. 62472064.

